# A Genome-wide Association and Admixture Mapping Study of Bronchodilator Drug Response in African Americans with Asthma

**DOI:** 10.1101/157198

**Authors:** Melissa L. Spear, Donglei Hu, Maria Pino-Yanes, Scott Huntsman, Anton S. M. Sonnenberg, Celeste Eng, Albert M. Levin, Marquitta J. White, Meghan E. McGarry, Neeta Thakur, Joshua M. Galanter, Angel C. Y. Mak, Sam S. Oh, Adam Davis, Rajesh Kumar, Harold J. Farber, Kelly Meade, Pedro C. Avila, Denise Serebrisky, Michael A. Lenoir, Emerita A. Brigino-Buenaventura, William Rodriquez Cintron, Shannon M. Thyne, Jose R. Rodriguez-Santana, Jean G. Ford, Rocio Chapela, Andrés Moreno Estrada, Karla Sandoval, Max A. Seibold, L. Keoki Williams, Cheryl A. Winkler, Ryan D. Hernandez, Dara G. Torgerson, Esteban G. Burchard

**Affiliations:** Department of Bioengineering and Therapeutic Sciences, University of California, San Francisco, San Francisco, CA, 94158; Department of Medicine, University of California, San Francisco, San Francisco, CA 94158; Research Unit, Hospital Universitario N.S. de Candelaria, Universidad de La Laguna, Tenerife, Spain, 38010; CIBER de Enfermedades Respiratorias, Instituto de Salud Carlos III, Madrid, Spain, 28029; Genomics and Health Group, Department of Biochemistry, Microbiology, Cell Biology and Genetics, Universidad de La Laguna (ULL), La Laguna (Tenerife), Spain, 38010; Department of Public Health Sciences, Henry Ford Health System, Detroit, MI; Department of Pediatrics, University of California, San Francisco, San Francisco, CA 94143; Department of Epidemiology and Biostatistics, University of California, San Francisco, San Francisco, CA, 94158; UCSF Benioff Children's Hospital Oakland, Center for Community Health and Engagement; Ann & Robert H. Lurie Children's Hospital of Chicago, Pediatrics, Chicago, Illinois, 60614; Department of Pediatrics, Section of Pulmonology, Baylor College of Medicine and Texas Children’s Hospital, Houston, TX, 77030; UCSF Benioff Children's Hospital Oakland, Oakland, California; Division of Allergy-Immunology, Northwestern University Feinberg School of Medicine, Chicago, Illinois, 60611; Pediatric Pulmonary Division, Jacobi Medical Center, Bronx, NY, 10461; Albert Einstein College of Medicine, Pediatrics, Bronx, New York; Bay Area Pediatrics, Oakland, CA, 94609; Department of Allergy & Immunology, Kaiser Permanente-Vallejo Medical Center, Vallejo, CA, 94589; Veterans Caribbean Health System, San Juan, Puerto Rico, 00921; Department of Pediatrics, David Geffen School of Medicine at ULCA, Olive View-UCLA Medical Center, Sylmar CA 91342; Centro de Neumologia Pediatrica, San Juan, Puerto Rico, 00917; Columbia University, New York, NY, 21205; Instituto Nacional de Enfermedades Respiratorias, Mexico City, MX, 14080; National Laboratory of Genomics for Biodiversity (LANGEBIO), CINVESTAV, Irapuato, Guanajuato 36821, Mexico; Department of Pediatrics, National Jewish Health, Denver, CO, 80206; Center for Health Policy and Health Services Research, Henry Ford Health System, Detroit, MI, 48202; Department of Internal Medicine, Henry Ford Health System, Detroit, MI, 48202; Basic Research Laboratory, National Cancer Institute, Leidos Biomedical Research, Frederick National Laboratory, Frederick, MD 21701; California Institute for Quantitative Biosciences (QB3), University of California, San Francisco, CA 94158; Institute for Human Genetics, University of California, San Francisco, CA, 94158

**Keywords:** genome-wide association study,, meta-analysis,, admixture mapping,, bronchodilator drug response,, albuterol,, African Americans,, Latinos,, asthma

## Abstract

**Background:** Short-acting B_2_-adrenergic receptor agonists (SABAs) are the most commonly prescribed asthma medications worldwide. Response to SABAs is measured as bronchodilator drug response (BDR), which varies among racial/ethnic groups in the U.S ^1, 2^. However, the genetic variation that contributes to BDR is largely undefined in African Americans with asthma^3^

**Objective:** To identify genetic variants that may contribute to differences in BDR in African Americans with asthma.

**Methods:** We performed a genome-wide association study of BDR in 949 African American children with asthma, genotyped with the Axiom World Array 4 (Affymetrix, Santa Clara, CA) followed by imputation using 1000 Genomes phase 3 genotypes. We used linear regression models adjusting for age, sex, body mass index and genetic ancestry to test for an association between BDR and genotype at single nucleotide polymorphisms (SNPs). To increase power and distinguish between shared vs. population-specific associations with BDR in children with asthma, we performed a meta-analysis across 949 African Americans and 1,830 Latinos (Total=2,779). Lastly, we performed genome-wide admixture mapping to identify regions whereby local African or European ancestry is associated with BDR in African Americans. Two additional populations of 416 Latinos and 1,325 African Americans were used to replicate significant associations.

**Results:** We identified a population-specific association with an intergenic SNP on chromosome 9q21 that was significantly associated with BDR (rs73650726, p=7.69 × 10^−9^). A trans-ethnic meta-analysis across African Americans and Latinos identified three additional SNPs within the intron of *PRKG1* that were significantly associated with BDR (rs7903366, rs7070958, and rs7081864, p≤5 × 10^−8^).

**Conclusions:** Our findings indicate that both population specific and shared genetic variation contributes to differences in BDR in minority children with asthma, and that the genetic underpinnings of BDR may differ between racial/ethnic groups.

**Key messages:**

- A GWAS for BDR in African American children with asthma identified an intergenic population specific variant at 9q21 to be associated with increased bronchodilator drug response (BDR).
- A meta-analysis of GWAS across African Americans and Latinos identified shared genetic variants at 10q21 in the intron of *PRKG1* to be associated with differences in BDR.
- Further genetic studies need to be performed in diverse populations to identify the full set of genetic variants that contribute to BDR.

## INTRODUCTION

Albuterol, a short-acting β_2_-adrenergic receptor agonist (SABA), is the most commonly prescribed asthma medication worldwide^4, 5^. SABAs cause rapid smooth muscle relaxation of the airways. Bronchodilator drug response (BDR) is a measure of a patient’s clinical response to SABA treatment and is quantitatively assessed as a change in forced expiratory volume in one second (FEV_1_) after administration of a SABA. BDR is a complex trait involving interactions among inflammatory cells^6^, airway epithelium^7^, smooth muscle cells^8^, and the autonomic nervous system^9^. Variation in BDR is likely influenced by both population-specific and shared environmental and genetic factors ^10-12^. In the United States (U.S.), BDR in children with asthma differs significantly between racial/ethnic groups^2, 10^. Specifically, African Americans have lower BDR compared to European populations even after controlling for asthma severity^13^. Compared to European Americans, African Americans suffer increased asthma morbidity and mortality ^2, 11, 14^ and decreased BDR likely contributes to these disparities in disease progression and outcomes. The extensive use of albuterol as a first-line therapy for asthma, coupled with the decreased drug response (BDR) and increased disease burden in African Americans underscores the importance of identifying genetic factors that influence BDR in African American children with asthma. Once identified, these factors may lead to the generation of novel therapies and targeted interventions that will serve to improve patient care and asthma outcomes in an overburdened and under-studied population.

To date, knowledge of genetic variation that contributes to BDR in African Americans is limited to a single genome-wide association study (GWAS) in 328 individuals^3^. Previous GWAS and candidate gene studies performed in populations of predominantly European ancestry with asthma have identified several BDR candidate genes^12, 15-24^. A recent study in Latinos with asthma replicated a number of these findings, and also identified novel population-specific associations with BDR^10^. Genetic effects identified in one population are not always generalizable across populations and several population-specific asthma-risk variants have been discovered in African-descent populations (i.e. African Americans and Latinos)^25-27^. Additionally, previous studies have shown that the varying degrees of African and European ancestry present in the African American population can be leveraged, through a technique known as admixture mapping, to identify the missing heritability of complex traits ^28^. Admixture mapping is a genome-wide approach that uses the variable allele frequencies of multiple SNPs between different ancestral populations to test for an association between local ancestry and phenotype 23, 25-27. The likelihood of population-specific effects, the limited number, and scale, of prior studies of BDR performed in African Americans, and ability to perform admixture mapping analysis highlights the possibility of gaining novel information through evaluating the impact of common genetic factors on BDR in African American children with asthma.

In this study, we performed a GWAS and admixture mapping study of bronchodilator drug response in 949 African American children with asthma from the Study of African Americans, Asthma, Genes & Environments (SAGE I and II)^29^. To increase power and distinguish between population-specific vs. shared associations, we also performed a trans-ethnic meta-analysis across our SAGE I and SAGE II participants and 1840 Latinos from GALA II (Genes-environments and Admixture in Latino Americans) studies^26^, respectively (total N=2,789). We further attempted replication of our population-specific and trans-ethnic metaanalysis results in 416 Latinos from the Genetics of Asthma in Latino Americans study (GALA I)^11, 30^ and 1,325 African Americans from the Study of Asthma Phenotypes and Pharmacogenomic Interactions by Race-Ethnicity (SAPPHIRE)^30, 31^.

## METHODS

### Study subjects from the Study of African Americans, Asthma, Genes & Environments

The Study of African Americans, Asthma, Genes & Environments (SAGE) is an ongoing case-control study of asthma in children and adolescents recruited from the San Francisco Bay Area in California^29^. Subjects were eligible if they were 8-21 years of age and self-identified all four grandparents as African American. Exclusion criteria included: (1) 10 or more pack-years of smoking; (2) any smoking within 1 year of recruitment date; (3) pregnancy in the third trimester; or (4) history of one of the following conditions: sickle cell disease, cystic fibrosis, sarcoidosis, cerebral palsy, or history of heart or chest surgery. Asthma was defined by physician diagnosis, asthma medication use and reported symptoms of coughing, wheezing, or shortness of breath in the 2 years preceding enrollment. Detailed clinical measurements were recorded for each individual whom DNA was collected from. In addition, trained interviewers administered questionnaires to obtain baseline demographic data, as well as information on general health, asthma status, social, and environmental exposures. Pulmonary function testing was conducted with a KoKo® PFT Spirometer (nSpire Health Inc., Louisville, CO) according to American Thoracic Society recommendations^32^, to obtain forced expiratory volume in one second (FEV_1_) in addition to other standard measurements of airway obstruction. Subjects with asthma were instructed to withhold their bronchodilator medications for at least 8 hours before testing. After completing baseline spirometry, subjects were given albuterol administered through a metered-dose inhaler (90 mcg/puff) with a spacer, and spirometry was repeated after 15 minutes to obtain post-bronchodilator measurements. The dose of albuterol was different in early stages of SAGE recruitment (2001-2005: SAGE I) than in more recent participants (2006-present: SAGE II). In SAGE I, post-bronchodilator FEV_1_ values were measured after providing the participants 2 puffs of albuterol (180 μg) if they were younger than 16 years of age and 4 puffs of albuterol (360 μg) if they were 16 years of age or older. In SAGE II, two doses of albuterol were delivered. For the first dose, 4 puffs of albuterol (360 μg) were provided independently of the age of the participant. For the second dose, two puffs (180ug) for children < 16 years old were administered and 4 puffs for subjects older ≥ 16 years.

Body mass index (BMI) was calculated for each participant using weight and height measures and converted to a categorical scale of underweight, normal, overweight, and obese according to the Centers for Disease Control and Prevention. For participants under 20 years old, standardized sex- and age-specific growth charts were used to calculate BMI percentiles (http://www.cdc.gov/nccdphp/dnpao/growthcharts/resources/sas.htm) and categorize their BMI as: underweight (BMI percentile<95^th^), normal (5^th^≤BMI<85^th^), overweight (85^th^≤BMI<95^th^), and obese (BMI≥95^th^). For participants older than 20 years old, BMI categories (http://www.cdc.gov/healthyweight/assessing/bmi/adultbmi/index.html-interpretedAdults) were defined as: underweight (BMI<18.5), normal (18.5≤BMI≤24.9), overweight (25≤BMI≤29.9) and obese (BMI≥30).

Institutional review boards approved the study and all subjects/parents provided written assent/consent, respectively.

### Genotyping and quality control (SAGE)

A total of 1,819 samples (1,011 asthma cases and 810 controls) were genotyped with the Axiom® World Array 4 (Affymetrix, Santa Clara, CA) at ~800,000 SNPs. Quality control was performed by removing SNPs that failed manufacturer’s quality control, had genotyping call rates below 95%, and/or had a deviation from Hardy-Weinberg equilibrium (p<10^−6^) within controls. 772,135 genotyped SNPs passed quality control. Samples were filtered based on discrepancy between genetic sex and reported gender and cryptic relatedness (PI_HAT>0.3). We excluded 3 subjects who were outliers for BDR (BDR of >60, or <-10). After sample quality control we included 759 SAGE II and 190 SAGE I asthma cases, for a total of 949 individuals with both genome-wide SNP data and measurements of bronchodilator drug response in the current study (Table 1).

**Table 1.** Descriptive statistics of SAGE I, SAGE II, & GALA II. Values shown are the means, with the standard deviation in parentheses.

|  | **SAGE I** | **SAGE II** | **GALA II** |
| --- | --- | --- | --- |
| **Total (N)** | 190 | 759 | 1830 |
| **Age (year)** | 18 (9.3) | 14 (3.6) | 13 (3.2) |
| **Sex (%Male)** | 41% | 52% | 55% |
| **Global African Ancestry** | 0.81 (0.13) | 0.72 (0.12) | 0.15 (0.13) |
| **Global Native American Ancestry** | - | - | 0.30 (0.25) |
| **BMI** |  |  |  |
| **<20 years** | 25 (7.3) (N=132) | 25 (7.2) (N=722) | 23(6.5) (N=1782) |
| **>20 years** | 31 (7.8) (N=58) | 29 (7.0) (N=37) | 30 (6.6) (N=40) |
| **Pulmonary Function** |  |  |  |
| **Pre-FEV1 % Predicted** | 92 (16) | 99 (14) | 91 (16) |
| **Pre-FVC % Predicted** | 100 (17) | 104 (13) | 95 (16) |
| **BDR** | 9 (9.1) | 9.5 (6.9) | 11 (8.2) |

Phasing of genotyped SNPs was performed using SHAPE-IT^33^, and genotype imputation was performed using IMPUTE2^34, 35^ using all populations from 1000 Genomes Project Phase III^36^ as a reference. Following imputation, a total of 9,573,507 genotyped and imputed (info score >0.3) SNPs with a MAF>0.05 were analyzed for SAGE II and 9,605,653 were analyzed for SAGE I.

### Study subjects from the Genes-environments & Admixture in Latino Americans study

A total of 1,830 Mexican and Puerto Rican asthma cases genotyped with the Axiom LAT1 array (World Array 4, Affymetrix) for the Genes-environments and Admixture in Latino Americans (GALA II) study were included in our analysis (Table 1 above). Specific details about the study are described elsewhere^10^. Briefly, imputation procedures identical to those described above for SAGE I and SAGE II were implemented, resulting in a total of 7,498,942 genotyped and imputed (info score >0.3) SNPs with a MAF>0.05.

### Study subjects from the Genetics of Asthma in Latino Americans study

For our replication phase, 247 Mexican and 169 Puerto Rican asthma cases genotyped with the Genome-Wide Human SNP Array 6.0 (Affymetrix) for the Genetics of Asthma in Latino Americans (GALA I) study were included. Details of the study are described elsewhere^11,30^.

### Study subjects from the Study of Asthma Phenotypes and Pharmacogenomic Interactions by Race-Ethnicity

For additional replication, we included 1,325 Africans Americans with asthma from the Study of Asthma Phenotypes and Pharmacogenomic Interactions by Race-Ethnicity (SAPPHIRE) genotyped with the Genome-Wide Human SNP Array 6.0 (Affymetrix). Estimates of local ancestry were obtained using RFMix^37^.Details of the study are described elsewhere ^25, 31^.

### Assessment of genetic ancestry

Genotypes from two populations were used to represent the ancestral haplotypes of African Americans for estimating local ancestry: HapMap European (CEU) and HapMap Africans (YRI). For Latinos, genotypes from 71 Native Americans were used as an additional ancestral population. Global ancestry was estimated using ADMIXTURE^38^ in a supervised analysis assuming two ancestral populations for African Americans and three ancestral populations for Latinos. A union set of SNPs was obtained by merging genotyped SNPs in SAGE and the ancestral populations (CEU/YRI). Local ancestry was estimated using the program LAMP-LD ^39^ in the GALA and SAGE studies and with RFMix in SAPPHIRE^37^.

### Genotype association testing

All statistical analyses were conducted using R (version 2.15.3). For SAGE individuals, we used standard linear regression to test for an association between BDR and allele dosage at each individual SNP, adjusting for age, sex, BMI category, and both global and local African ancestry. A GWAS of BDR in GALA II has been previously published^10^, however, this previous work did not include adjustment for BMI; in this study we re-ran the GWAS using a new reference imputation panel and further adjusted by BMI^40^. For GALA II individuals, we adjusted for age, sex, BMI category, ethnicity, global Native American and African ancestry, and local ancestry. All analyses were performed using imputed genotypes from 1000 Genomes phase III. Using the fixed-effects model implemented in METAL^41^, we performed a meta-analysis of common variants (MAF ≥ 5%) across African Americans (SAGE I and SAGE II) and Latinos (GALA II). We selected variants that were common (MAF ≥ 5%) within each individual study and then took the intersection of SNPs for the meta-analysis.

### Admixture mapping

We used local ancestry estimates generated across the genome to perform admixture mapping in African Americans. Linear regression models adjusted for age, sex, BMI category, and global African ancestry were used to identify significant associations between local ancestry estimates and BDR. The threshold for genome-wide significance was calculated using the empirical autoregression framework with the package *coda* in R to estimate the total number of ancestral blocks^42, 43^. The Bonferroni threshold was calculated as α=2.4 × 10^−4^ based on 245 ancestral blocks. Admixture mapping was performed separately in SAGE I and SAGE II and combined in a meta-analysis using METAL^41^.

### Replication in GALA I and SAPPHIRE

We attempted replication of significant population-specific (SAGE I and SAGE II) and cosmopolitan (SAGE I, SAGE II, GALA II) associations with BDR in the GALA I and SAPPHIRE studies. Replication in GALA I was performed using genotype imputation (i.e. *in silico* replication), followed by an examination at a locus-wide level for SNPs within +/- 50 kb. We imputed 100 kb regions around each SNP using the program IMPUTE2 for Mexican and Puerto Rican participants run separately using 1000 Genomes phase 3 haplotypes as a reference. Linear regression was used to test foran association between allele dosage and BDR separately in Mexicans and Puerto Ricans, adjusting for age, sex, BMI category, global and local ancestry. Replication in SAPPHIRE was also performed using linear regression to test for an association between allele dosage and BDR in African Americans while adjusting for age, sex, BMI category, and global and local African ancestry. For GALA I and SAPPHIRE replication, statistical significance at the SNP level was evaluated at p<0.05, and at the locus-wide level was established using a conservative Bonferroni correction adjusting by the number of SNPs within +/- 50 kb of the original candidate SNP.

## RESULTS

### Genome-wide association results

After filtering variants with a MAF ≥ 5% and with imputation quality score (info score) ≥ 0.3, we tested for an association of BDR at a total of 9,190,349 SNPs in 949 African Americans with asthma (λ *=* 1.006). We identified a single genome-wide significantly associated SNP within an intergenic region on chromosome 9 (rs73650726, imputation quality score=0.86) (Figures 1, 2, 3, and Table 2 below). At this variant, additional copies of the A1 allele (A), was associated with decreased drug response (β=-3.8, p=7.69 × 10^−9^) (Table 2 below).

**Figure 1:**
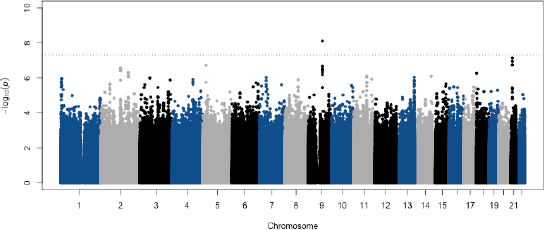
Meta-analysis of genome-wide association studies with BDR in African. Americans. Association testing for BDR was performed using linear regression including age, sex, BMI category, local and global ancestry as covariates separately in SAGE I and II and combined in a meta-analysis. Dotted line indicates the genome-wide significance threshold of 5 × 10^−8^.

**Figure 2:**
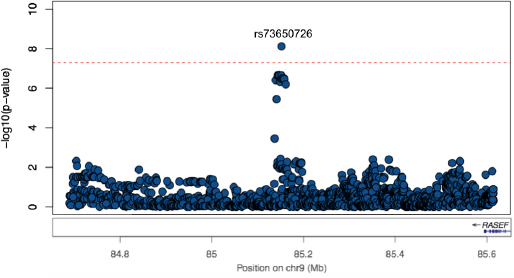
LocusZoom plot of chr9:84653000-85653000. Region includes genotyped and imputed variants from 1000 Genomes phase 3. Blue = variants common in SAGE I and II. Dotted line indicates the genome-wide significance threshold of 5 × 10^−8^.

**Figure 3:**
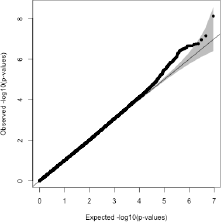
Quantile-quantile plots for genome-wide allelic associations with BDR in a meta-analysis of. SAGE I and II (inflation factor: λ = 1.006 for ~10 M common SNPs)

**Table II:** Genome-wide significant associations identified through a meta-analysis within African Americans (SAGE I and II), and within African Americans and Latinos (SAGE I, SAGE II, and GALA II). Under ‘Direction’ the first symbol refers to SAGE I, second to SAGE II,and third to GALA II. 0 = absent/rare in study

| **African Americans (SAGE I and II):** |  |  |  |  |  |  |  |  |
| --- | --- | --- | --- | --- | --- | --- | --- | --- |
| Chr | SNP | Position (hg19) | A1 | A2 | Effect (A1) | StdErr | Pvalue | Direction |
| 9q21 | rs73650726 | 85152666 | A | G | -3.8 | 0.66 | $7.69 \times 10^{-9}$ | --0 |
| **African Americans + Latinos (SAGE I, SAGE II, GALA II):** |  |  |  |  |  |  |  |  |
| Chr | SNP | Position (hg19) | A1 | A2 | Effect (A1) | StdErr | Pvalue | Direction |
| 10q21 | rs7903366 | 53689774 | T | C | 1.23 | 0.22 | $3.94 \times 10^{-8}$ | +++ |
| 10q21 | rs7070958 | 53691116 | A | G | -1.24 | 0.23 | $4.09 \times 10^{-8}$ | --- |
| 10q21 | rs7081864 | 53690331 | A | G | 1.23 | 0.22 | $4.94 \times 10^{-8}$ | +++ |

The SNP rs73650726 is common in African Americans but rare in Latinos, with a minor allele frequency of 8% in both SAGE studies, but at a frequency of 1% in GALA II (Table III below). This is consistent with allele frequencies observed in the 1000 Genomes Project, where the variant is common in African populations (8%), rare in Latino populations (1-2%), and absent in European and Asian populations (Figure 4 below)^44^.

**Table III:** Individual genome-wide association study results from SAGE I, SAGE II, and GALA II.

| **SAGE I** |  |  |  |  |  |  |  |  |
| --- | --- | --- | --- | --- | --- | --- | --- | --- |
| **Chr** | **SNP** | **Position (hg19)** | **A1** | **A2** | **Freq (A1)** | **Effect (A1)** | **StdErr** | **Pvalue** |
| 9q21 | rs73650726 | 85152666 | A | G | 0.92 | -2.99 | 1.92 | 0.12 |
| 10q21 | rs7903366 | 53689774 | C | T | 0.4 | -0.54 | 1.02 | 0.6 |
| 10q21 | rs7070958 | 53691116 | A | G | 0.39 | -0.5 | 1.03 | 0.63 |
| 10q21 | rs7081864 | 53690331 | G | A | 0.39 | -0.46 | 1.03 | 0.65 |

| **SAGE II** |  |  |  |  |  |  |  |  |
| --- | --- | --- | --- | --- | --- | --- | --- | --- |
| **Chr** | **SNP** | **Position (hg19)** | **A1** | **A2** | **Freq (A1)** | **Effect (A1)** | **StdErr** | **Pvalue** |
| 9q21 | rs73650726 | 85152666 | A | G | 0.92 | -3.9 | 0.7 | $3.36 \times 10^{-8}$ |
| 10q21 | rs7903366 | 53689774 | C | T | 0.41 | -1.66 | 0.39 | $2.83 \times 10^{-5}$ |
| 10q21 | rs7070958 | 53691116 | A | G | 0.41 | -1.66 | 0.4 | $3.22 \times 10^{-5}$ |
| 10q21 | rs7081864 | 53690331 | G | A | 0.41 | -1.65 | 0.4 | $3.26 \times 10^{-5}$ |

| **GALA II** |  |  |  |  |  |  |  |  |
| --- | --- | --- | --- | --- | --- | --- | --- | --- |
| **Chr** | **SNP** | **Position (hg19)** | **A1** | **A2** | **Freq (A1)** | **Effect (A1)** | **StdErr** | **Pvalue** |
| 9q21 | rs73650726 | 85152666 | A | G | 0.99 | 0.29 | 1.25 | 0.81 |
| 10q21 | rs7903366 | 53689774 | C | T | 0.6 | -1.06 | 0.28 | $1.74 \times 10^{-4}$ |
| 10q21 | rs7070958 | 53691116 | A | G | 0.6 | -1.08 | 0.28 | $1.57 \times 10^{-4}$ |
| 10q21 | rs7081864 | 53690331 | G | A | 0.6 | -1.06 | 0.28 | $1.79 \times 10^{-4}$ |

**Figure 4:**
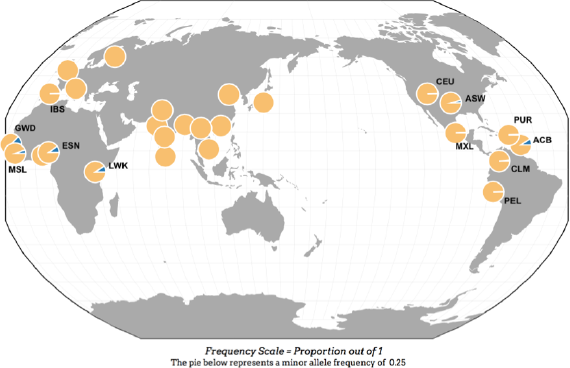
Geographic distribution of allele frequencies of rs73650726. Each pie chart refers to a population from the 1000 Genomes Project phase 3. Yellow= Major allele (A), blue = minor allele ÍGV rs73650726 is common onlv in DODulations with African ancestry.

In order to increase power and identify associations shared between populations we performed a trans-ethnic meta-analysis across African American, Mexican, and Puerto Rican participants from SAGE I, SAGE II, and GALA II, respectively. Following quality control and filtering on variants common in each study (MAF ≥ 5%), we took the overlap between the three studies and we performed a meta-analysis on 6,570,864 SNPs (λ = 1.004). We identified genome-wide significant associations at three SNPs located on chromosome 10 within the intron of *PRKG1*: rs7903366 (β=1.23, p=3.94 × 10^−8^), rs7070958 (β=-1.24, p=4.09 × 10^−8^), and rs7081864 (β=1.23, p=4.94 × 10^−8^) (imputation quality scores > 0.98, Figures 5, 6 & 7 below, Tables II & III above). All three SNPs are eQTLs for *PRKG1* in lung tissue from the GTEx database (Table IV below)^45^, with the minor allele associated with decreased expression.

**Figure 5:**
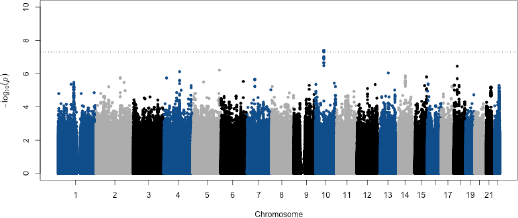
Meta-analysis of genome-wide association studies with BDR in African Americans and Latinos. Association testing for BDR was performed using linear regression including age, sex, BMI category, local and global ancestry as covariates; including ethnicity for GALA II. Dotted line indicates the genome-wide significance threshold of 5 × 10^−8^.

**Figure 6:**
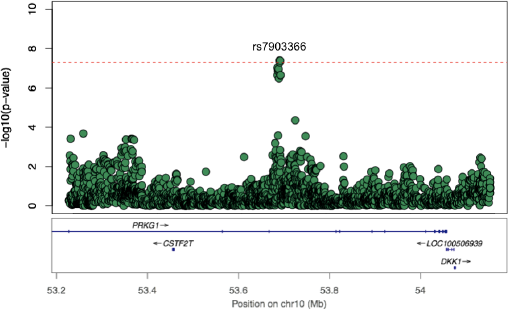
LocusZoom plot of chr10:53200000-54200000. Region includes genotyped and imputed variants from 1000 Genomes phase 3. Green = variants common in SAGE I, SAGE IIand GALA II. Dotted line indicates the genome-wide significance threshold of 5 × 10^−8^.

**Figure 7:**
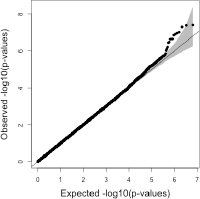
Quantile-quantile plots for genome-wide allelic associations with BDR in a metaanalysis of. SAGE I, SAGE II, and GALA II (inflation factor: λ = 1.004 for all common SNPs)

**Table IV:** Correlation between the expression of *PRKG1* in the lung and minor alleles at three intronic SNPs associated with BDR (cis-eQTLs). Data is from the GTEx database.

| SNP | Reference Allele | Pvalue | Effect Size | T-Statistic | StdErr | Tissue | Gene |
| --- | --- | --- | --- | --- | --- | --- | --- |
| rs7903366 | C | 0.00051 | -0.12 | -3.5 | 0.034 | Lung | PRKG1 |
| rs7070958 | A | 0.00046 | -0.12 | -3.6 | 0.034 | Lung | PRKG1 |
| rs7081864 | G | 0.00052 | -0.12 | -3.5 | 0.034 | Lung | PRKG1 |

Replication of African American population-specific (rs73650726) and shared (rs7903366, rs7070958, rs7081864) was attempted in two independent Latino (GALA I) and African American (SAPPHIRE) studies. Although none of the identified associations replicated in either study population, the African American population-specific association between rs73650726 and BDR, identified in the SAGE studies, displayed a similar trend in direction of effect in Puerto Ricans (GALA I; β = -6.2) and in African Americans (SAPPHIRE, β = -0.65) (Table V below). In addition, none of the SNPs within 50 kb of the four original SNPs were significantly associated with BDR following Bonferroni correction (Table VI below). Lastly, we evaluated previously identified candidate SNPs from prior candidate gene and GWAS with BDR in patients with asthma. After accounting for fifteen comparisons, no SNPs met the statistical significance threshold (p<3.33 × 10^−3^) (Table VII below); only rs9551086 in *SPATA13* had a p-value below 0.05 (p=0.02).

**Table V:** Replication results of candidate SNPs in GALA I and SAPPHIRE.

| **GALA I** |  |  |  |  |  |  |  |  |
| --- | --- | --- | --- | --- | --- | --- | --- | --- |
| **Population** | **Chr** | **SNP** | **Position (hg19)** | **A1** | **A2** | **Effect (A1)** | **StdErr** | **Pvalue** |
| Mexicans | 9q21 | rs73650726 | 85152666 | A | G | 1.71 | 10.24 | 0.87 |
| Mexicans | 10q21 | rs7903366 | 53689774 | C | T | 1.02 | 0.99 | 0.30 |
| Mexicans | 10q21 | rs7070958 | 53691116 | A | G | 1.04 | 0.99 | 0.29 |
| Mexicans | 10q21 | rs7081864 | 53690331 | G | A | 1.03 | 0.99 | 0.30 |
| Puerto Ricans | 9q21 | rs73650726 | 85152666 | A | G | -6.22 | 3.87 | 0.11 |
| Puerto Ricans | 10q21 | rs7903366 | 53689774 | C | T | -0.51 | 1.01 | 0.61 |
| Puerto Ricans | 10q21 | rs7070958 | 53691116 | A | G | -0.53 | 1.00 | 0.60 |
| Puerto Ricans | 10q21 | rs7081864 | 53690331 | G | A | -0.53 | 1.00 | 0.60 |

**Table V: Replication results of candidate SNPs in GALA I and SAPPHIRE.**
| **SAPPHIRE** |  |  |  |  |  |  |  |
| --- | --- | --- | --- | --- | --- | --- | --- |
| **Chr** | **SNP** | **Position (hg19)** | **A1** | **A2** | **Effect (A1)** | **StdErr** | **Pvalue** |
| 9q21 | rs73650726 | 85152666 | A | G | -0.65 | 0.94 | 0.49 |
| 10q21 | rs7903366 | 53689774 | C | T | 0.91 | 0.55 | 0.10 |
| 10q21 | rs7070958 | 53691116 | A | G | 0.87 | 0.55 | 0.11 |
| 10q21 | rs7081864 | 53690331 | G | A | 0.84 | 0.55 | 0.13 |

**Table VI:** Top replication results (MAF >0.05) +/-50 kb of candidate SNPs in GALA I and SAPPHIRE.

| **GALA I** |  |  |  |  |  |  |  |  |  |
| --- | --- | --- | --- | --- | --- | --- | --- | --- | --- |
| **Population** | **Chr** | **SNP** | **Position (hg19)** | **A1** | **A2** | **Freq (A1)** | **Effect (A1)** | **StdErr** | **Pvalue** |
| Mexicans | 9 | rs34293766 | 85126730 | T | G | 0.94 | 5.74 | 2.27 | 0.01 |
| Mexicans | 10 | rs59960792 | 53690251 | C | T | 0.80 | 2.97 | 1.12 | 0.01 |
| Puerto Ricans | 9 | rs10780548 | 85167648 | A | G | 0.63 | 2.93 | 0.99 | 0.003 |
| Puerto Ricans | 10 | rs6480581 | 53672743 | T | C | 0.74 | -2.67 | 1.08 | 0.01 |

| **SAPPHIRE** |  |  |  |  |  |  |  |  |  |
| --- | --- | --- | --- | --- | --- | --- | --- | --- | --- |
| **Population** | **Chr** | **SNP** | **Position (hg19)** | **A1** | **A2** | **Freq (A1)** | **Effect (A1)** | **StdErr** | **Pvalue** |
| African Americans | 9 | rs62576848 | 85144677 | T | C | 0.94 | 3.03 | 1.23 | 0.01 |
| African Americans | 10 | rs35969600 | 53662118 | C | T | 0.72 | -1.49 | 0.59 | 0.01 |

**Table VII.** Candidate SNP replication in meta-analysis of SAGE I and SAGE II. Under ‘Direction’ column, 1^st^ symbols refer to SAGE I, second refer to SAGE II.

| **GENE** | **SNP** | **Chr** | **Position** | **A1** | **A2** | **Effect** | **Pvalue** | **Direction** |
| --- | --- | --- | --- | --- | --- | --- | --- | --- |
| *SOCS-ASB3* region | rs350729<sup>23</sup> | 2 | 52983773 | T | G | 0.22 | 0.53 | -+ |
| *ADRB2* | rs1042713<sup>15</sup> | 5 | 148206440 | A | G | -0.13 | 0.7 | -- |
| *ADRB2* | rs1042714<sup>15</sup> | 5 | 148206473 | C | G | -0.56 | 0.22 | -- |
| *ADCY9* | rs2230739<sup>18</sup> | 16 | 4033436 | T | C | -0.14 | 0.77 | +- |
| *CRHR2* | rs7793837<sup>17</sup> | 7 | 30726777 | A | T | -0.26 | 0.58 | -- |
| *ARG1* | rs2781659<sup>19</sup> | 6 | 131891820 | A | G | 0.39 | 0.3 | -+ |
| *SPATS2L* | rs295137<sup>24</sup> | 2 | 201150040 | T | C | 0.4 | 0.23 | -+ |
| *SPATS2L* | rs295114<sup>24</sup> | 2 | 201195602 | T | C | 0.41 | 0.22 | -+ |
| *THRB* | rs892940<sup>20</sup> | 3 | 24538838 | A | G | 0.52 | 0.21 | ++ |
| *CRHR2* | rs73294475<sup>10</sup> | 7 | 30701596 | T | C | -0.32 | 0.51 | +- |
| *SPATA13* | rs9507294<sup>3</sup> | 13 | 24823347 | T | C | 0.38 | 0.48 | ++ |
| *SPATA13* | rs912142<sup>3</sup> | 13 | 24827500 | A | G | -0.23 | 0.51 | -- |
| *SPATA13* | rs2248119<sup>3</sup> | 13 | 24827094 | A | G | 0.49 | 0.16 | ++ |
| *SPATA13* | rs9551086<sup>3</sup> | 13 | 24830330 | T | C | -1.36 | 0.02 | -- |
| *SPATA13* | rs9553225<sup>3</sup> | 13 | 24823006 | A | G | -1.17 | 0.10 | -- |

### Admixture mapping results

We tested for an association of BDR with local genetic ancestry inferred at 478,441 SNPs in 949 African Americans with asthma (190 from SAGE I and 759 from SAGE II) (Figures 8 & 9 below). A meta-analysis across both studies yielded no significant associations with ancestry (p<2.4 × 10^−4^) (Figure 10 below). The most significant peak was located on chromosome 8p11 where African ancestry was associated with higher BDR (β=1.49, p=6.34 × 10^−4^) (Table VIII below).

**Figure 8:**
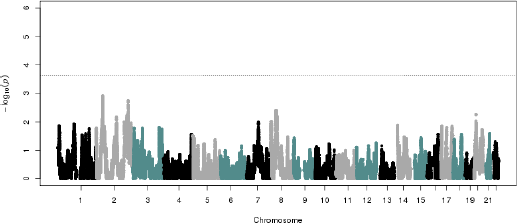
Admixture mapping in SAGE II (n= 759) for African ancestry and BDR. Ancestry association testing was performed at 478,441 markers using linear regression including age, sex, BMI category, and global African ancestry covariates.

**Figure 9:**
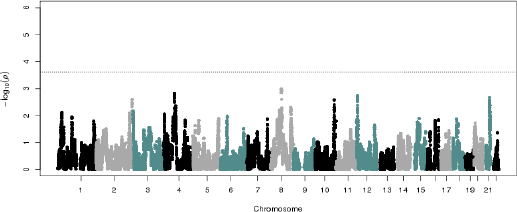
Admixture mapping in SAGE I (n= 190) for African ancestry and BDR. Ancestry association testing was performed at 478,441 markers using linear regression including age, sex, BMI category, and global African ancestry covariates.

**Figure 10:**
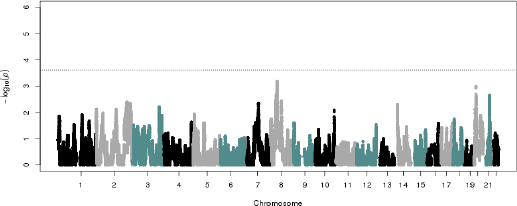
Meta-analysis of admixture mapping in SAGE I and II (n=949) for African ancestry and BDR. Ancestry association testing was performed at 478,441 markers using linear regression including age, sex, BMI category, and global African ancestry covariates.

**Table VIII:** Meta-analysis of admixture mapping results within African Americans (SAGE I and II). Under ‘Direction’ the first symbol refers to SAGE I, second to SAGE II

| Chr | SNP | Position (hg19) | Effect | StdErr | Pvalue | Direction |
| --- | --- | --- | --- | --- | --- | --- |
| 8p11 | rs3927941 | 39805797 | 1.49 | 0.44 | $6.34 \times 10^{-4}$ | ++ |

## DISCUSSION

We performed a genome-wide association study for bronchodilator drug response in African Americans, and identified a population-specific association between rs73650726, located on chromosome 9, and BDR. Specifically, we discovered that the G allele of rs73650726 was associated with increased BDR and is more common in African Americans compared to European populations (Table II above). The variant rs73650726, located on chromosome 9, does not map to any gene, but SNPs in high linkage disequilibrium (r^2^≥0.8) with this marker are located in enhancer histone marks in lung tissues ^46^.

Our results demonstrate that population-specific genetic variation contributes to variation in BDR in African American children with asthma. We further combined our results in a metaanalysis for BDR in African Americans and Latinos and identified multiple intronic variants in *PRKG1* that were associated with BDR in both populations. Overall our results demonstrate that population-specific and shared genetic factors contribute to variation in BDR among African American children with asthma. Through investigating the underlying genetics of BDR in diverse populations we have uncovered novel information regarding both the pathophysiology and pharmacogenetics of asthma.

Three of our significantly associated variants fell within the intronic region of an annotated gene, Protein Kinase, CGMP-Dependent, Type I *(PRKG1). PRKG1* encodes for a cyclic GMP-dependent protein kinase, which phosphorylates proteins involved in regulating platelet activation and adhesion^47^, gene expression^48, 49^, vascular smooth muscle cell contraction^50^, and feedback of the nitric-oxide (NO) signaling pathway^51^. Notably, the NO pathway is a key pathway in modulating vasodilation in response to beta-agonists such as albuterol via beta 2-adrenergic receptors^52^, making *PRKG1* a highly plausible BDR candidate gene. The three SNPs are in high linkage disequilibrium (r^2^≥0.8) with variants known to be functional^46^, and are all associated with the expression of *PRKG1* in the lung - a tissue highly relevant to BDR. From the Genotype-Tissue Expression (GTEx) project database, the reference allele for all three SNPs was associated with decreased expression of the gene in lung tissue^45^. Thus, additional studies are required to identify the causal underlying variation at this locus, such as direct sequencing of this locus, and how the expression of *PRKG1* may be related to differences in BDR.

We attempted to replicate our study findings and candidate SNPs previously found to be associated with BDR, however we found no significant associations following multiple testing corrections. This could be due to differences in study design, the presence of population specific differences in genetic contributions to BDR, and/or varying patterns of linkage disequilibrium between populations. Furthermore, we were limited in sample size in GALA I^25^ to evaluate associations at low frequency variants, and note that SAPPHIRE is comprised of mainly adults^31^ in comparison to SAGE and GALA II that are comprised of mainly children.

In conclusion, we identified two novel loci with biological plausibility whereby genetic variation is associated with differential response to albuterol, the most commonly prescribed asthma medication. One of these loci contains variation associated with BDR that is common to African Americans, a population that has historically been understudied in genetic studies^53-55^. Further genetic studies in African Americans is essential for identifying a more comprehensive set of genetic variants that contribute to differences in BDR, which in turn will lead to a better understanding of the pharmacogenetic response to asthma therapies. This will provide the foundation for genetic risk profiling and precision medicine, identifying novel genes and pathways associated with BDR, and the development of novel asthma therapies.

## Acknowledgments

The authors acknowledge the patients, families, recruiters, health care providers, and community clinics for their participation in SAGE and GALA II. In particular, we thank study coordinator Sandra Salazar and the recruiters who obtained the data: Duanny Alva, MD; Gaby Ayala-Rodriguez; Lisa Caine; Elizabeth Castellanos; Jaime Colon; Denise DeJesus; Blanca Lopez; Brenda Lopez, MD; Louis Martos; Vivian Medina; Juana Olivo; Mario Peralta; Esther Pomares, MD; Jihan Quraishi; Johanna Rodriguez; Shahdad Saeedi; Dean Soto; and Ana Taveras.

The contents of this publication are solely the responsibility of the authors and do not necessarily represent the official views of the NIH.

## Abbreviations used

BDR: bronchodilator drug response
BMI: body mass Index
FEV_1_: forced expiratory volume in one second
GALA I: Genetics of Asthma in Latino Americans
GALA II: Genes-environments & Admixture in Latino Americans
GWAS: genome-wide association study
MAF: minor allele frequency
SABA: short-acting β_2_-agonist
SAGE: Study of African Americans, Asthma, Genes, & Environments
SAPPHIRE: Study of Asthma Phenotypes and Pharmacogenomic Interactions by Race-Ethnicity
SNP: single nucleotide polymorphism

